# Understanding the molecular basis for enhanced glutenase activity of actinidin

**DOI:** 10.1101/2023.05.24.542047

**Authors:** Shivangi Puja, Shreya Seth, Rachna Hora, Satinder Kaur, Prakash Chandra Mishra

## Abstract

Management of gluten intolerance is currently possible only by consumption of gluten free diet(GFD) for a lifetime. The scientific community has been searching for alternatives to GFD, like inclusion of natural proteases with meals or pre-treatment of gluten containing foods with glutenases. Actinidin from kiwifruit has shown considerable promise in digesting immunogenic gliadin peptides as compared to other plant derived cysteine proteases. Through this article, we have attempted to understand the structural basis for elevated protease action of actinidin against gliadin peptides by using an *in silico* approach. Docking experiments reveal key differences between the binding of gliadin peptide to actinidin and papain, which may be responsible for their differential digestive action. Sequence comparison of different plant cysteine proteases highlights amino acid residues surrounding the active site pocket of actinidin that are unique to this molecule and hence likely to contribute to its digestive properties.

**Graphical summary:** 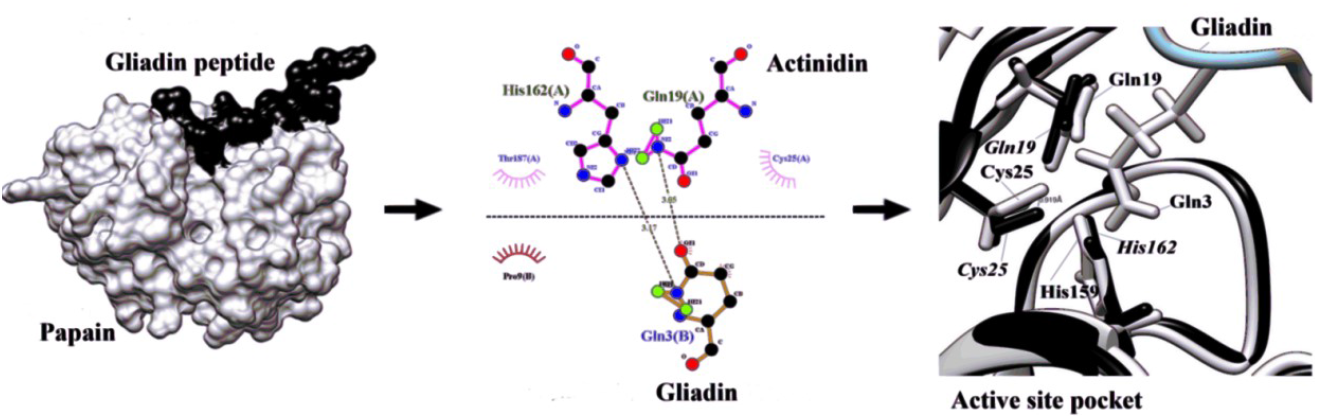

## INTRODUCTION

Gluten related disorders are a growing health issue worldwide^1^. These manifest either as the autoimmune celiac disease (CD) in genetically predisposed individuals, or wheat allergy (WA) or non-celiac gluten sensitivity (NCGS). Though gastrointestinal symptoms like diarrhea *etc*. for these diseases overlap, the underlying reasons are discrete. CD is defined by the presence of circulating autoantibodies against tissue transglutaminase (tTG) resulting in enteropathy and sometimes extraintestinal problems like anemia, osteoporosis and abnormal liver function^2^.Occurrence of CD is strongly associated with the presence of HLA DQ2 and DQ8 genes in affected individuals. According to serologic test results and biopsydata,1.4% and 0.7% people have CD respectively, which is believed to be a gross underestimate owing to unreported asymptomatic and mildly symptomatic cases. WA is characterized by the presence of IgE class of antibodies against gliadin, a component of gluten. NCGS is also an immune mediated disorder that occurs more commonly than CD and WA (up to 13% of the population being affected^3^), and is characterized by absence of autoantibodies and lesions in the duodenal mucosa. All the above manifestations of gluten intolerance are provoked by gluten protein’s highly immunogenic gliadin component.

Glutens form a storage protein family present in wheat, barley, rye and some oat cultivars^4^. Wheat gluten contains gliadins and glutenins which provide elasticity and softness to the dough. Homologues of glutens are present in barley (hordeins), rye (secalins) and oats (avenins), which have the ability to trigger gluten related disorders. All these glutens are rich in amino acids prolines (P) and glutamines (Q), making them refractory to digestion by pepsin or other pancreatic enzymes which fail to work on P containing peptide bonds^5^.These partially digested P and Q rich peptides from gluten act as epitopes that evoke an inflammatory response in celiac disease patients resulting in atrophy of the intestinal villi, crypt hyperplasia and nutrient malabsorption. Shan *et al* had reported a 33 mer peptide from wheat alpha gliadin to be the most immunogenic^6^.Currently, treatment of individuals suffering from any form of gluten intolerance is a strict gluten-freediet (GFD)^7^ which is extremely difficult to follow and may have compromised nutrient composition. The scientific community has been long seeking alternatives to GFD including usage of P and Q specific gluten degrading enzymes. These enzymes that facilitate digestion of immunogenic peptides have shown promise as therapeutic agents for gluten intolerance.

Glutenases may be used to pre-treat flours in order to render them hypoallergenic or they can be consumed alongside a meal to act as digestive aids^8^.Most of the enzymes used in this approach are prolyl endopeptidases (PEP) of bacterial origin i.e., from *Flavobacterium, Sphingomonas* and *Myxococcus* species. However, since their pH optima lies between 7-8, they are rapidly inactivated under gastric conditions in the presence of pepsin. Also, a fungal PEP from *Aspergillus niger* which works best at pH 4-5 was unable to sufficiently degrade gluten in three separate phase I clinical trials. Therefore, exploitation of alternate glutenases, preferably from plant-based natural sources is the need of the hour^9^. Several cysteine proteases from plants have shown potential for digesting gliadin and its peptides^10^. These include papain from *Carica papaya*, triticains from germinating wheat, bromelain from *Ananas comosus*, actinidin from *Actinidia chinesis etc*. Of these, actinidin has shown notable potential for gluten degradation.

Actinidin, a kiwifruit cysteine protease has been shown to improve digestion of a variety of dietary proteins using a two stage *in vitro* digestion system^11^. This study had highlighted the enhanced ability of actinidin to efficiently cleave whey protein isolate, zein, gluten, and gliadin. Consumption of kiwifruit also improved the rate at which digested nitrogen entered the small intestine (SI) and the apparent amino acid digestibility at proximal and medial SI in pigs fed with a beef based diet^12^. Therefore, actinidin can facilitate digestion of protein-rich foods, supporting the case for kiwifruit consumption as a digestive aid. Another study compared the potential of actinidin and other cysteine proteases from fruits to digest isolated glutens and digestion resistant gliadin peptides under simulated gastrointestinal conditions^5^. Actinidin could efficiently cleave gluten proteins, the extremely resistant and immunogenic 33-mergliadin peptide and peptides comprising QQQ/PFP found in gluten epitopes. Actinidin showed better apparent degree of hydrolysis while digesting gliadin peptides as compared to papain, bromelain and two other commercial glutenases sourced from *Aspergillus* species. Their LC-MS/MS data displayed that actinidin could cleave numerous peptide bonds at the N-and C-termini of P residues under gastric conditions. In the present study, we have followed an *in silico* approach to understand the molecular basis for the enhanced ability of actinidin to cleave immunogenic gliadin peptides. We have used molecular docking and multiple sequence alignment to gain new insights on structural aspects of actinidin that may be responsible for its augmented activity. Our findings may form the basis for development of newer proteases by protein engineering that are effective in digestion of immunogenic gluten peptides.

## MATERIALS AND METHODS

### 1. Sequence and structure retrieval

Sequences for plant cysteine proteases with glutenase activity were retrieved from UniProt (http://www.uniprot.org)^13^. Coordinates of three dimensional structures of papain (1BP4)^14^, actinidin (1AEC)^15^ and gliadin peptide P31-43 (6QAX)^16^ were retrieved from RCSB protein data bank for docking, visualization and structure analysis. Multiple sequence alignment of the retrieved plant cysteine protease sequences was performed by using CLUSTAL omega^17^.

### 2. Molecular docking of cysteine proteases with gliadin peptide and structural analysis

Coordinates of the three dimensional structure of gliadin peptide P31-43 (6QAX) (LGQQQPFPPQQPY)were used for its docking with papain (1BP4). Docking experiment was carried out by HADDOCK^18^, which is a flexible docking program that is information driven. The best docked complex i.e. the top cluster which is considered to be the most reliable by HADDOCK was used for further analysis. The crystallographic structure of actinidin (1AEC) was merged with the papain - gliadin docked structure using UCSF Chimera^19^. Binding energy and residues involved in protein - protein interaction were studied by PRODIGY^20^, LigPlot^+^ tool^21^ and PyMol^22^. PRODIGY predicts binding energy between two proteins based on their intermolecular contacts. LigPlot^+^ is used fortwo dimensional plotting of residue-residue interactions between a protein and its interacting partner via an intuitive java interface. PyMOL, a python-based software, is a visualization system which is used for visualization and analysis.

## RESULTS AND DISCUSSION

### Structural analysis highlights distinct interactions in the active site pockets of papain and actinidin

Papain is a broad specificity cysteine protease that cleaves after basic amino acids, leucine and glycine^23^ but it was found to be less efficient than actinidin at digesting gliadin peptides. The structure of papain (1BP4) was used for docking with the gliadin peptide P31-43 (6QAX) using HADDOCK, and the best complex generated was selected for analysis. The immunogenic peptide bound at the active site pocket of papain (Fig 1a). The active site of papain lies within a V-shaped cleft at the interface of two subdomains and hosts the catalytic diad (Cys25 - His159 ion pair)^14^. While Asn175 and Gln19 are pertinent for positioning of the catalytic diad, Trp177 is responsible for generation of its nucleophilic character. Our binding pocket analysis of the papain – gliadin P31-43 pair shows hydrogen bond formation between Gln3 of gliadin peptide and Cys25, His159 and Gln19 of papain (Fig 1b). Visualization of the same in the complex structure clearly shows these hydrogen bonds (Fig 1c). Residues of papain involved in hydrophobic interactions are Cys22, Gly23, Cys63, Gln142, Leu143, Trp177 and Gly180 (Fig 1b).

**Figure 1:**
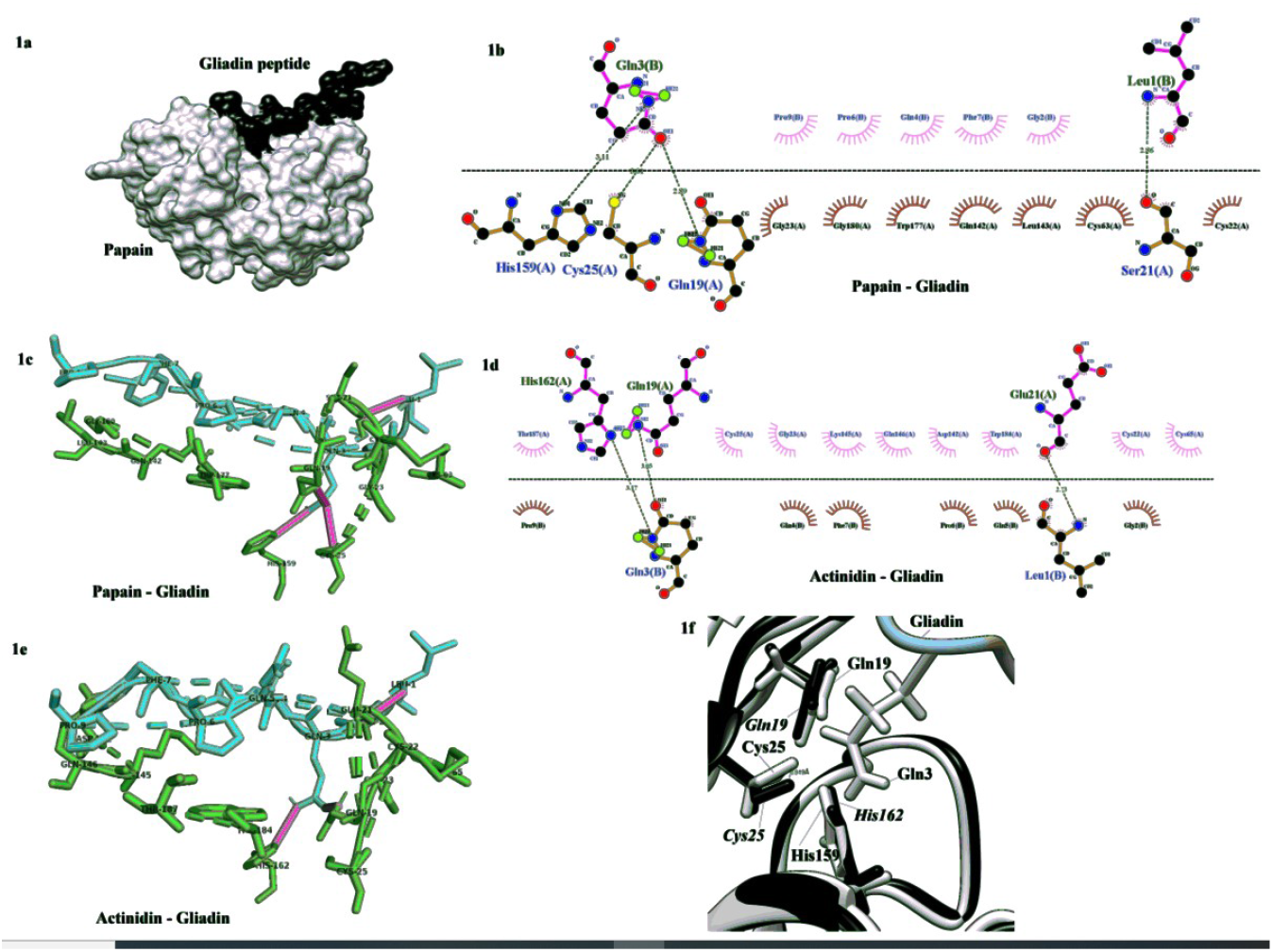
Papain/ actinidin–gliadin interaction. 1a: Docked structure of papain (grey) (1BP4) with gliadin peptide P31-43 (black) (6QAX) using HADDOCK showing the peptide bound at the active site pocket of papain. The model shown was generated by using UCSF Chimera. 1b:LigpPlot^+^ analysis showing residue-residue interaction between papain and gliadin peptide. Horizontal dashed line shows the interface (top: gliadin; bottom: papain). Green dotted lines display hydrogen bonds while arced residues depict hydrophobic interactions. 1c: Visualization of hydrogen bonds (pink) between Gln3 of gliadin peptide (cyan) and Cys25, His159 and Gln19 of papain(green) using PyMol.1d:Ligplot^+^analysis showing interfacial contacts between actinidin (top) and gliadin peptide (bottom). Interface, Hydrogen bonds and hydrophobic interactions are shown as explained in Fig 1b. 1e: Figure showing hydrogen bonds (pink)formed by Gln3 of gliadin (cyan) and Gln19 & His162of actinidin (green).1f: Merged structure showing shift of 0.919 Åin Cys25 of actinidin vs Cys25 of papain.

Gliadin peptide P31-43 includes the QQQ/PFP sequence that is reported to be more efficiently cleaved by actinidin as compared to papain^5^. Actinidin is a papain-like cysteine protease from kiwifruit whose catalytic diad is formed by Cys25and His162^24^. Notably, actinidin is reported to act as a proline specific endopeptidase^5^ showcasing its ability to digest P rich peptides. In order to compare the catalytic pockets of papain and actinidin, we overlapped the structure of actinidin (1AEC)on the papain-gliadin complex (merged structure). In comparison to papain, Cys25 of actinidin was unable to form a hydrogen bond with Gln3 of the gliadin peptide (Fig 1d and 1e) while Gln 19 and His 162 were engaged in hydrogen bond formation. Amino acids of actinidin facilitating hydrophobic interactions with the gliadin peptide were Cys22, Gly23, Cys25, Cys65, Asp142, Lys145, Gln146, Trp184 and Thr187 (Fig 1d). Interestingly, Gln5 of the gliadin peptide was able to form hydrophobic interactions with actinidin residues but not with papain (Figs 1b & 1d). This residue lies at the cleavage site QQQ/PFP. Further analysis of the merged structure revealed that active site Cys25 of actinidin was shifted with respect to Cys25 of papain by 0.919 Å, possibly disallowing it to form a hydrogen bond with the peptide Gln3 and altering its catalytic properties (Fig 1f). However, the active site histidines and glutamines 19 of both the structures overlapped well. The binding energy of the actinidin–peptide complex was also estimated to be lower (−9.2 kCal/mol, Kd = 1.8E-07) as compared to the papain–peptide pair (−8.8 kCal/mol, Kd = 3.7E-07), suggesting a relatively stronger interaction of actinidin with the gliadin peptide. While interfacial contacts by apolar amino acids dominated in the papain-gliadin peptide pair, contribution of polar and charged residues in contact formation was significantly more in the actinidin–gliadin peptide complex (Table S1).

### Sequence analysis of plant cysteine proteases reveals unique residues in actinidin

*Actinidia chinensis* and *deliciosa* are two cultivars of kiwifruit that are sourced from China (Chinese gooseberry) and New Zealand (golden kiwifruit) respectively^25,26^. The sequences of actinidins from *A. chinensis* and *deliciosa* were found to be 98% identical with variations at residues 118 (Q to R), 193 (V to G), 226 (V to L), 279 (I to T), 352 (S to P) and 366 (N to K) (Figure S1). All these alterations were found to lie far away from the active site regions of the enzyme.

In order to understand the differences in biomolecular interactions and binding energies of papain and actinidin with gliadin peptide, sequence analysis of active site regions of plant cysteine proteases was performed. Multiple sequence alignment of full length sequences of actinidin, papain, bromelain, five distinct cysteine proteases from germinating wheat (triticain alpha, triticain beta, triticain gamma & two other cysteine peptidases) and EP-B2 (from germinating barley) revealed that the active site region of all these proteases are conserved (Figure 2). The catalytic region of actinidin lies within the amino acid stretches 145 - 156, 286-296, and 303-322^15^(Figure S2). Most residues within the catalytic regions of the aligned cysteine proteases are either strictly conserved, or show significant variation across different enzymes. However, two residues within these regions were found to be conserved in all the compared enzyme sequences except actinidin. All proteases except actinidin contain a Serine (Ser) residue at position 150 that is adjacent to the active site Cys151 (corresponding to Cys25 of the crystallographic structures). This is replaced by a Glyresidue (highlighted in grey and underlined) in actinidin. Also, a conserved glycine residue in all the aligned cysteine proteases is exchanged for an aspartic acid (Asp) at position 311 (highlighted in grey and underlined). This Asp residue lies at the immediate carboxy end of Trp310 (same as Trp184 of actinidin and corresponding to Trp177 of papain),which has a critical role in building the nucleophilic character of the catalytic diad. These identified residues surrounding the active site pocket of actinidin are likely to have an effect on intermolecular interactions and binding energy as noticed in the above docking experiment. Testing the enzymatic activity and specificity of site directed mutants for these sites in actinidin would confirm the role of these residues in enzyme action.

**Figure 2.**
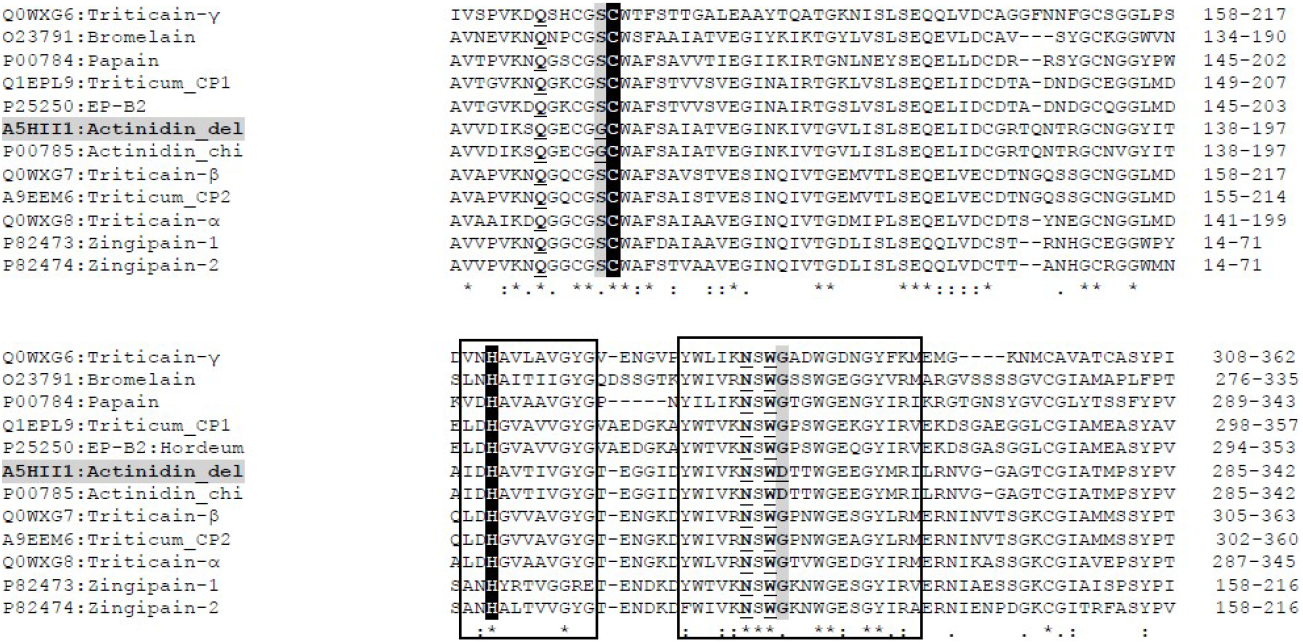
Multiple sequence alignment of plant cysteine proteases. The catalytic regions are boxed. Active site catalytic diad Cys-His ion pair is highlighted in black. Residues known to influence the catalytic diad i.e. Q145, N308 and W310 in actinidin along with their counterparts in other cysteine proteases are underlined. Residues G150 and D311 of actinidin identified through our analysis are highlighted in grey and underlined, while their corresponding S and G amino acids from other proteases are highlighted in grey.

## Conclusion

Actinidin has an exceptional quality among natural fruit proteases to efficiently digest immunogenic gliadin peptides. It works well at concentrations above 2.7 U/mL and at pH values greater than 2. This concentration of actinidin equates consumption of two kiwifruits in a meal^27^. Usage of this protease may be developed as an alternative treatment to gluten free diet. Understanding the structural basis for the activity and specificity of actinidin is likely to pave the path for its engineering and hence development of therapeutics against diseases arising from gluten intolerance.

## Abbreviations

GFD: gluten free diet;
CD: celiac disease;
PEP: prolyl endopeptidase;
WA: wheat allergy;
NCGS: non celiac gluten sensitivity

## ETHICS APPROVAL AND CONSENT TO PARTICIPATE

Not applicable.

## HUMAN AND ANIMAL RIGHTS

No animals/humans were used for studies that are the basis of this research.

## CONSENT FOR PUBLICATION

Not applicable.

## AVAILABILITY OF DATA AND MATERIALS

Not applicable.

## ACKNOWLEDGEMENTS

Declared none.

---

**Figure S1:**
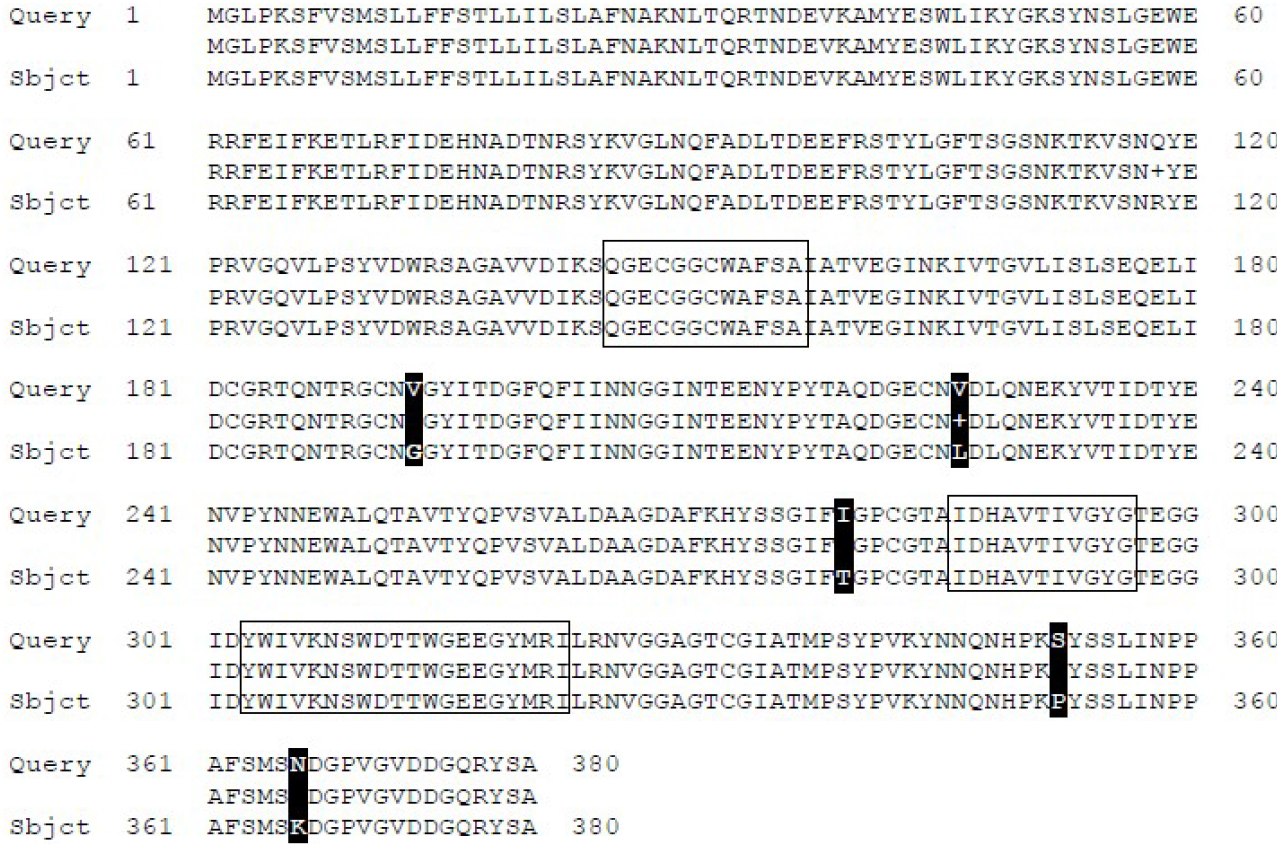
Pairwise alignment using BLAST2 for sequences of actinidins from *A. chinensis* (query) and *deliciosa* (sbjct) showing 98% identity. Sequence variations are highlighted in black. Active site regions are boxed.

**Figure S2:**
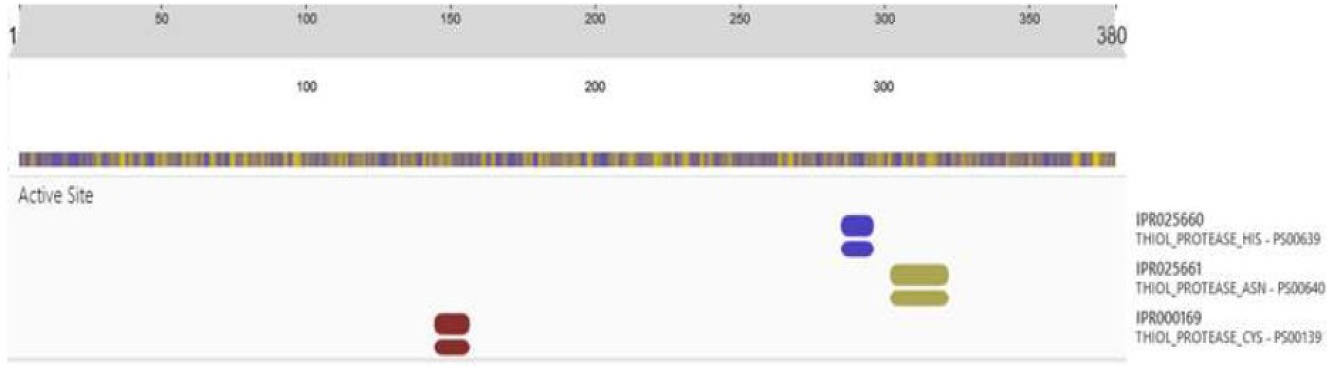
Result of domain analysis of actinidin sequence using InterPro.

**Table S1:** Predicted values for bindingenergies (ΔG) and dissociation constants (Kd) f**or** papain-gliadin and actinidin**-**gliadin pairs are shown. The number of interfacial contacts formed in each pair are tabulated according to their properties.

| Parameters | Papain | Actinidin |
| --- | --- | --- |
| $\Delta G$ (kcal mol<sup>-1</sup>) | -8.8 | -9.2 |
| $K_d$ (M) at 25.0 °C | 3.70E-07 | 1.80E-07 |
| **Number of Interfacial Contacts (ICs) per property** |  |  |
| ICs charged-charged | 0 | 0 |
| ICs charged-polar | 2 | 5 |
| ICs charged-apolar | 1 | 5 |
| ICs polar-polar | 5 | 1 |
| ICs polar-apolar | 17 | 14 |
| ICs apolar-apolar | 14 | 10 |
| **Non Interacting Surface (NIS) per property** |  |  |
| NIS charged | 19.35% | 19.76% |
| NIS apolar | 40.00% | 40.12% |

## REFERENCES

(1) Balakireva, A. V.; Zamyatnin Jr, A. A. Properties of Gluten Intolerance: Gluten Structure, Evolution, Pathogenicity and Detoxification Capabilities. Nutrients2016, 8 (10), 644.

(2) Calado, J.; Machado, M. V. Celiac Disease Revisited. GE-Port. J. Gastroenterol.2022, 29 (2), 108–121.

(3) Czaja-Bulsa, G. Non Coeliac Gluten Sensitivity–A New Disease with Gluten Intolerance. Clin. Nutr.2015, 34 (2), 189–194.

(4) Levy, J.; Bernstein, L.; Silber, N. Celiac Disease: An Immune Dysregulation Syndrome. Curr. Probl. Pediatr. Adolesc. Health Care2014, 44 (11), 324–327.

(5) Jayawardana, I. A.; Boland, M. J.; Higgs, K.; Zou, M.; Loo, T.; Mcnabb, W. C.; Montoya, C. A. The Kiwifruit Enzyme Actinidin Enhances the Hydrolysis of Gluten Proteins during Simulated Gastrointestinal Digestion. Food Chem.2021, 341, 128239.

(6) Shan, L.; Molberg, Ø.; Parrot, I.; Hausch, F.; Filiz, F.; Gray, G. M.; Sollid, L. M.; Khosla, C. Structural Basis for Gluten Intolerance in Celiac Sprue. Science2002, 297 (5590), 2275–2279.

(7) Niewinski, M. M. Advances in Celiac Disease and Gluten-Free Diet. J. Am. Diet. Assoc.2008, 108 (4), 661–672.

(8) Wei, G.; Helmerhorst, E. J.; Darwish, G.; Blumenkranz, G.; Schuppan, D. Gluten Degrading Enzymes for Treatment of Celiac Disease. Nutrients2020, 12 (7), 2095.

(9) Jayawardana, I. A.; Montoya, C. A.; McNabb, W. C.; Boland, M. J. Possibility of Minimizing Gluten Intolerance by Co-Consumption of Some Fruits–A Case for Positive Food Synergy? Trends Food Sci. Technol.2019, 94, 91–97.

(10) Savvateeva, L. V.; Gorokhovets, N. V.; Makarov, V. A.; Serebryakova, M. V.; Solovyev, A. G.; Morozov, S. Y.; Reddy, V. P.; Zernii, E. Y.; Zamyatnin Jr, A. A.; Aliev, G. Glutenase and Collagenase Activities of Wheat Cysteine Protease Triticain-α: Feasibility for Enzymatic Therapy Assays. Int. J. Biochem. Cell Biol.2015, 62, 115–124.

(11) Kaur, L.; Rutherfurd, S. M.; Moughan, P. J.; Drummond, L.; Boland, M. J. Actinidin Enhances Protein Digestion in the Small Intestine as Assessed Using an in Vitro Digestion Model. J. Agric. Food Chem.2010, 58 (8), 5074–5080.

(12) Montoya, C. A.; Cabrera, D. L.; Zou, M.; Boland, M. J.; Moughan, P. J. The Rate at Which Digested Protein Enters the Small Intestine Modulates the Rate of Amino Acid Digestibility throughout the Small Intestine of Growing Pigs. J. Nutr.2018, 148 (11), 1743–1750.

(13) Apweiler, R.; Bairoch, A.; Wu, C. H.; Barker, W. C.; Boeckmann, B.; Ferro, S.; Gasteiger, E.; Huang, H.; Lopez, R.; Magrane, M. UniProt: The Universal Protein Knowledgebase. Nucleic Acids Res.2004, 32 (Suppl_1), D115–D119.

(14) LaLonde, J. M.; Zhao, B.; Smith, W. W.; Janson, C. A.; DesJarlais, R. L.; Tomaszek, T. A.; Carr, T. J.; Thompson, S. K.; Oh, H.-J.; Yamashita, D. S. Use of Papain as a Model for the Structure-Based Design of Cathepsin K Inhibitors: Crystal Structures of Two Papain-Inhibitor Complexes Demonstrate Binding to S ‘-Subsites. J. Med. Chem.1998, 41 (23), 4567–4576.

(15) Varughese, K. I.; Su, Y.; Cromwell, D.; Hasnain, S.; Xuong, N. H. Crystal Structure of an Actinidin-E-64 Complex. Biochemistry1992, 31 (22), 5172–5176.

(16) Calvanese, L.; Nanayakkara, M.; Aitoro, R.; Sanseverino, M.; Tornesello, A. L.; Falcigno, L.; D’Auria, G.; Barone, M. V. Structural Insights on P31-43, a Gliadin Peptide Able to Promote an Innate but Not an Adaptive Response in Celiac Disease. J. Pept. Sci.2019, 25 (5), e3161.

(17) Sievers, F.; Higgins, D. G. Clustal Omega for Making Accurate Alignments of Many Protein Sequences. Protein Sci.2018, 27 (1), 135–145.

(18) De Vries, S. J.; Van Dijk, M.; Bonvin, A. M. The HADDOCK Web Server for Data-Driven Biomolecular Docking. Nat. Protoc.2010, 5 (5), 883–897.

(19) Pettersen, E. F.; Goddard, T. D.; Huang, C. C.; Couch, G. S.; Greenblatt, D. M.; Meng, E. C.; Ferrin, T. E. UCSF Chimera—a Visualization System for Exploratory Research and Analysis. J. Comput. Chem.2004, 25 (13), 1605–1612.

(20) Xue, L. C.; Rodrigues, J. P.; Kastritis, P. L.; Bonvin, A. M.; Vangone, A. PRODIGY: A Web Server for Predicting the Binding Affinity of Protein–Protein Complexes. Bioinformatics2016, 32 (23), 3676–3678.

(21) Wallace, A. C.; Laskowski, R. A.; Thornton, J. M. LIGPLOT: A Program to Generate Schematic Diagrams of Protein-Ligand Interactions. Protein Eng. Des. Sel.1995, 8 (2), 127–134.

(22) Yuan, S.; Chan, H. S.; Hu, Z. Using PyMOL as a Platform for Computational Drug Design. Wiley Interdiscip. Rev. Comput. Mol. Sci.2017, 7 (2), e1298.

(23) Haertlé, T. Enzymes: Analysis and Food Processing. In Encyclopedia of Food and Health; Caballero, B., Finglas, P. M., Toldrá, F., Eds.; Academic Press: Oxford, 2016; pp 524–531. https://doi.org/10.1016/B978-0-12-384947-2.00257-9.

(24) Novinec, M.; Lenarčič, B. Papain-like Peptidases: Structure, Function, and Evolution. Biomol. Concepts2013, 4 (3), 287–308.

(25) He, X.; Fang, J.; Chen, X.; Zhao, Z.; Li, Y.; Meng, Y.; Huang, L. Actinidia Chinensis Planch.: A Review of Chemistry and Pharmacology. Front. Pharmacol.2019, 10, 1236.

(26) Shastri, K. V.; Bhatia, V.; Parikh, P. R.; Chaphekar, V. N. Actinidia Deliciosa: A Review. Int. J. Pharm. Sci. Res.2012, 3 (10), 3543.

(27) Montoya, C. A.; Rutherfurd, S. M.; Olson, T. D.; Purba, A. S.; Drummond, L. N.; Boland, M. J.; Moughan, P. J. Actinidin from Kiwifruit (Actinidia Deliciosa Cv. Hayward) Increases the Digestion and Rate of Gastric Emptying of Meat Proteins in the Growing Pig. Br. J. Nutr.2014, 111 (6), 957–967.

